# Transcriptional contingency in the early hematopoiesis of inherited bone marrow failure syndromes

**DOI:** 10.64898/2026.11.02.763843

**Authors:** Angel A. De la Cruz, Pablo Siliceo, Ulises Juárez-Figueroa, Sara Frías, Paula Leal-Anaya, Alfredo Rodríguez

**Author notes:** **Corresponding Author:** Alfredo Rodríguez, PhD, Bone Marrow Failure & Carcinogenesis Laboratory, National Institute of Pediatrics (INP), Department of Genomic Medicine & Environmental Toxicology, Institute of Biomedical Research, Universidad Nacional Autónoma de México (UNAM), Av. Universidad 3000, Universidad Nacional Autónoma de México, C.U. Delegación Coyoacán, C.P. 04510, Mexico City.

## Abstract

The inherited bone marrow failure syndromes (IBMFS) are a group of heterogeneous genetic diseases caused by germline pathogenic variants in genes involved in hematopoiesis and genomic integrity. Despite sharing common phenotypical manifestations that have aided opportune diagnosis, such as bone marrow failure, congenital malformations and an increased predisposition to develop cancer, little is known about the shared and distinctive transcriptional programs in their hematopoietic stem and progenitor cells compartment that underlie the clinical manifestations of each disease.

In this work we have successfully integrated and analyzed the publicly available scRNAseq datasets of HSPCs from patients with Fanconi anemia (FA) and Telomere Biology disorders (TBD), as well as healthy controls. Once integrated, we performed quality control metrics, as well as batch effect correction using different methods. Through dimensionality reduction, cell clustering and cell type annotation we found distinctive gene expression profiles per syndrome and per cell type, as well as orchestrated gene expression profiles, including upregulation of inflammatory pathways and cell proliferation pathways.

This work shows the integration of datasets from FA and TBD, two major IBMFS, and provides the basis for an accumulative, comparative, and integrated atlas of scRNAseq data from patients with IBMFS.

## INTRODUCTION

Hematopoiesis, the process by which blood cells are formed, depends on the appropriate function of the hematopoietic stem and progenitor cells (HSPCs). HSPCs are pluripotent cells residing in the bone marrow (BM) with the capacity to produce lymphomyeloid progenitors and megakaryocyte-erythroid progenitors, which in turn will give rise to all the mature blood components. Hematopoiesis is a rigorous and fundamental process involving various critical pathways essential for maintaining efficient blood production (1).

A malfunctioning BM becomes evident when a patient presents with aplastic anemia (AA), defined by the presence of peripheral blood cytopenias accompanied by a hypoplastic bone marrow, a condition known as bone marrow failure (BMF) (2,3). Etiological subgroups of BMF have been delineated to facilitate clinical management, these are categorized primarily into acquired BMF and inherited BMF, which account for approximately 80-85% and 15-20% of cases in the pediatric context, respectively (4–9). Inherited defects in the regulation of HSPCs, such as those affecting telomere maintenance, DNA damage repair or ribosomal function, can lead to BMF (2,3,8).

Inherited bone marrow failure syndromes (IBMFS) are a group of monogenic disorders that present with hematological abnormalities in the form of AA and predisposition to myelodysplastic neoplasm (MDS), acute myeloid leukemia (AML), and increased risk to develop solid tumors. These syndromes might also present extra-hematological abnormalities, including congenital malformations (1,8,10).

The most common, and better known, IBMFS are dyskeratosis congenita (DC), a telomere biology disorder (TBD), and Fanconi anemia (FA), a DNA repair disorder (10). Although different in their etiology, both IBMFs are fundamentally underlined by a disrupted capacity to maintain the integrity of their genome structure, either by telomere uncapping in TBDs, or by increased chromosome breakage in FA. TBDs are commonly diagnosed through the telomeric length measurement via Flow-FISH and FA by using the diepoxybutane (DEB) chromosome breakage test (11,12).

TBDs encompass a spectrum of syndromic affections that can be detected in the adulthood as AA, MDS, AML, pulmonary fibrosis or cryptogenic hepatic cirrhosis, or during childhood, when DC is diagnosed through a mucocutaneus triad of nail dystrophy, oral leukoplakia and reticulated skin pigmentation. To date, 17 genes have been associated with TBDs (*DKC1, TERT, TERC, TINF2, TCAB1, NOP10, NHP2, CTC1, RTEL1, TPP1, PARN, POT1, NAF1,* and *STN1*) (13,14). Patients with FA typically present with a multisystemic phenotype, involving multiple birth defects, the most notorious include cardiac abnormalities, renal abnormalities, and radial ray defects. FA is due to the inheritance of biallelic pathogenic variants in genes that participate in the FA/BRCA pathway of DNA repair (*FANCA, FANCB, FANCC, FANCD1, FANCE, FANCF, FANCG, FANCI, FANCJ, FANCL, FANCM, FANCN, FANCO, FANCP, FANCR, FANCS, FANCT, FANCV, FANCW*). Both TBDs and FA present genetic heterogeneity (15,16).

BMF is a leading cause of early mortality in patients with FA and TBD, who typically develop BMF during childhood, whereas in some patients, especially in those with a TBD, BMF might not appear but until adulthood. Hematopoietic stem cell transplantation (HSCT) from a healthy donor remains the only curative option for the hematologic complications observed in these patients, however other comorbidities remain (8,17,18).

Recent advancements in single-cell RNA sequencing (scRNAseq) have begun to uncover the complex cellular heterogeneity within bone marrow populations, offering a glimpse into the molecular disruptions that contribute to BMF (19). While applying scRNA-seq to patients with TBDs and FA can reveal both commonalities and unique disruptions in HSPCs function, many of the underlying pathways driving BMF remain poorly understood. Continued research is essential to fully elucidate these unknown mechanisms (20). Identifying novel molecular pathways and potential biomarkers through scRNAseq analysis could pave the way for more targeted and personalized therapeutic approaches, ultimately improving outcomes for patients with IBMFS.

In this work we have integrated and analyzed publicly available scRNAseq datasets of Lin^-^ CD34^+^ cells of BM samples from patients with FA, TBDs, and healthy controls. We aimed to identify transcriptional contingency and perform comparative functional characterization of scRNAseq datasets of HSPC from patients with IBMFS. Through dimensionality reduction, cell clustering and cell type annotation we have found distinctive clusters with shared cell types between syndromes as well as populations with unique transcriptional programs who differed between syndromes. With this comparative work we are starting to find, not only the expected divergencies in the transcriptional profiles of these IBMFS, but also commonalities that upon proper scrutiny might uncover shared pathological mechanism and even guide the development of shared therapies.

## METHODS

### Data download

We downloaded the pre-processed scRNA-Seq data from the public repository Gene Expression Omnibus (GEO), encompassing human HSPCs from 6 patients with FA (GSE157591) (35) and 4 patients with TBDs (GSE169709) (36); as well as 9 healthy donors, 5 from GSE157591 and 4 from GSE169709, respectively.

### Data processing and quality control

The downloaded count matrices were transferred to the R statistical environment (v4.2.0) for further processing with Seurat (v5.1.0) (39); only intersecting genes between both datasets (GSE157591 and GSE169709) were considered for downstream analysis. Data assessment showed that standard Seurat quality control metrics were already implemented on FA samples (nFeature_RNA > 200, nFeature_RNA < 2500, percent.mt < 5), allowing for no additional filtering. In TBD samples, cells with more than >25% of gene expression coming from mitochondrial genes (percent.mt < 25) where filtered out, avoiding substantial cell dropout when standard 5% was implemented. Finally, all data was normalized with a standard log1p transformation with the Seurat function NormalizeData (*normalization.method = LogNormalize*).

### Data integration and dimensionality reduction

All samples were concatenated into a single composite Seurat object. Data integration was conducted by implementing different data integration methods to assess optimal batch effect correction between datasets (RPCAIntegration, FastMNNIntegration, HarmonyIntegration, scVIIntegration). Integration benchmark scores assessing the performance of batch effect removal were calculated in all integration methods using Seurat’s native function MixingMetric (k = 5, dims 1:2, max.k = 300).

FastMNNintegration integration results were used for downstream processing based on overall performance scores. After integration, 10 principal dimensions from FastMNNintegration anchored on 2000 highly variable genes calculated by the “vst” method with Seurat function FindVariableFeatures (s*election.method = ‘vst’, nfeatures = 2000*), were used for all UMAP embeddings.

### Clustering and cell annotation

Unsupervised clustering of the data was made using nearest-neighbor graph construction under the Seurat function FindNeighbors (*k.param = 20*). Different resolutions were scanned for cluster determination using Seurat’s function FindClusters (resolution = 0.1,0.2,0.4,0.6,0.8,1,1.1). A resolution of 1.1 was chosen for cluster determination given that it produced the highest number of clusters without group redundancy.

Cell annotation was made in a hybrid manner, manually annotating cell types using canonical cell markers and differentially expressed genes among clusters, as well as referencing automatic annotations obtained through Azimuths Bone Marrow reference dataset (v.1)(32).

### Differential gene expression analysis

Differentially expressed genes between groups were calculated through the FindAllMArkers() function in Seurat using a Wilcoxon test and filtering genes shown in the results with a Log2FC > 1. The expression of top DEGs was plotted for each cell type and group as violin plots for further comparison.

### Cell communication analysis

Inference of cell-cell communication was done using CellChat(v1.5)(31). The processed datasets were divided converted to a CellChat object using createCellChat(). Using the human ligand-receptor CellChat database (31) as reference, under the category “secreted signaling”, overly expressed genes related to secreted cell signaling and cell interaction were subset from each dataset by means of the CellChat functions identifyOverExpressedGenes() and identifyOverExpressedInteractions(), respectively.

Cell communication networks at a signaling pathway level were constructed by calculating communication probabilities between cell types, excluding detected communications between groups with less than 5 cells, using the computeCommunProbPathway() function. The compositional network resuming all interactions for each group was obtained with the function aggregateNet().

Altered interactions between biological conditions and cell types were identified by comparing the number of interactions and their respective strength from the inferred communication networks using the comparedInteractions() function.

## RESULTS

### Integration of scRNA-Seq data from HSPCs from FA and TBDs shows heterogeneous cell type composition between syndromes

We integrated publicly available scRNAseq datasets comprising HSPCs from 6 patients with FA (GSE157591) and 4 with TBDs (GSE169709), as well as 9 healthy donors, 5 and 4 from each dataset respectively. Both datasets underwent specific quality control filtering to retain high quality cells only (**Figure 1A**). Despite being generated by the same sequencing technology (10x), UMAP embeddings showed visible batch effect between datasets and individuals (**Fig 2A**). To attain reliable integration of the data without diluting biological variation, different integration methods were implemented on the data (RPCAIntegration, FastMNNIntegration, HarmonyIntegration, scVIIntegration). By benchmarking a mixing score (MixingMetric()) across methods accounting for batch correction, we identified FastMNNintegration and RPCAintegration as the integration tools with the highest mixing scores.

**Figure 1.**
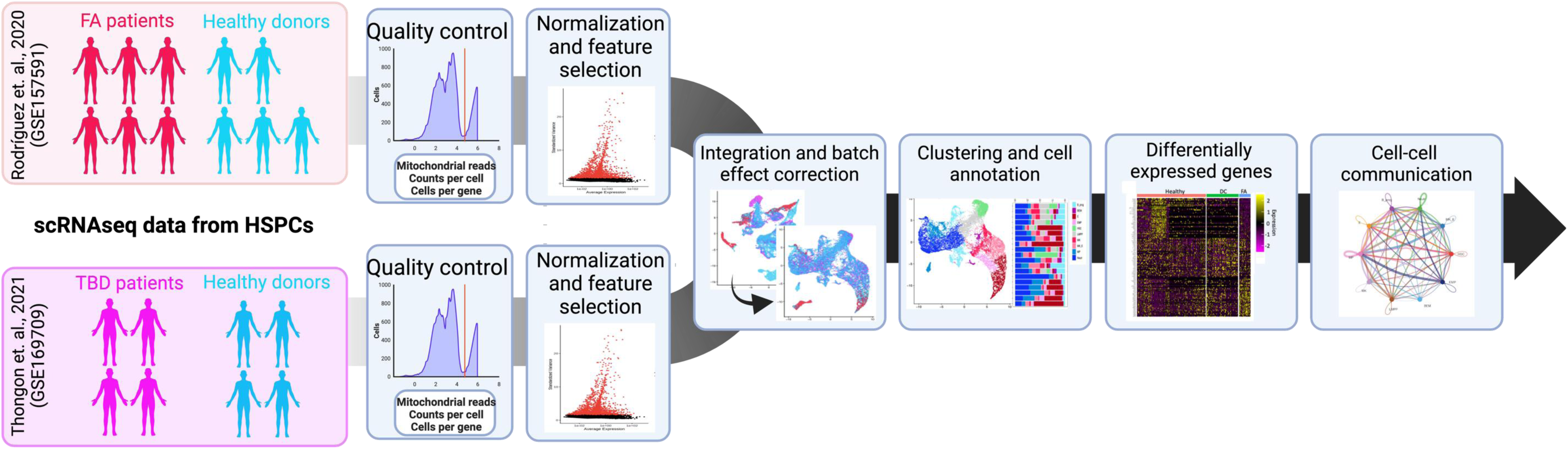
General workflow of scRNASeq IBMFS data processing, integration and analysis. scRNAseq datasets from HSCPs of patients with FA and TBD, along with healthy donors, were downloaded from the GEO omnibus data repository (reference number shown). Datasets underwent cell and gene filtering based on standard quality control metrics. Normalization of the data and selection of variable features (2000) were used for data integration and batch correction, benchmarking different integration tools. Data analysis consisted in clustering, cell annotation, calculation of differentially expressed genes and inference of cell-cell communication based on ligand-receptor pairings.

**Figure 2.**
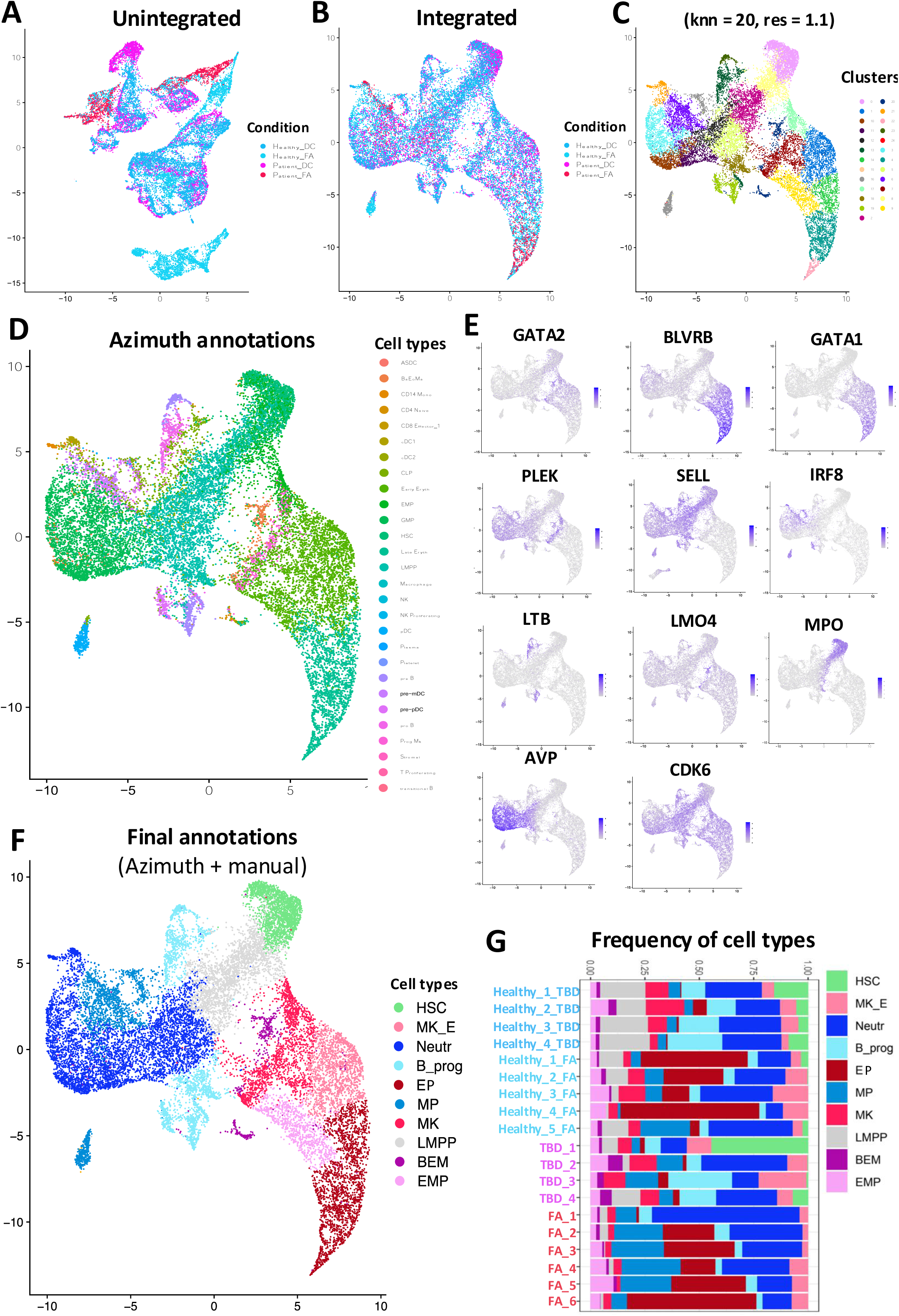
Data integration, clustering and cell annotation of FA and DC scRNAseq datasets. **(A)** UMAP displaying the data before integration, two healthy datasets were included, as well as one dataset from patients with FA and one dataset from patients with DC. **(B)** UMAP of the integrated data using the FastMNNintegration tool. **(C)** UMAP showing data clustering from the integrated data using a *K* nearest neigbour value of 20 and clustering resolution of 1.1. **(D)** Data annotation using the Azimuth mask algorithm. **(E)** UMAPs displaying the expression of canonical gene markers for cell types of interest. **(F)** UMAP showing final annotations resulting from combining manual and Azimuth. **(G)** Proportions of annotated cell types across samples from healthy donors and patients.

FastMNNintegration was chosen for downstream embeddings (**Figure 2B**) contrary to RPCAintegration, due to a finer cluster definition when unsupervised clustering was performed, yielding a total of 25 clusters (**Fig 2C**). Cell annotation using Azimuth resulted in a total of 28 distinctive cell types which included lineage positive cell annotations like macrophages, effector CD8+ T cells, dendritic cells, among others (**Fig 2D**). Considering that Azimuth’s mask detection is based on transcriptional similarities with previously annotated cell types, overestimation of cell types in the samples was bypassed by coupling automatic annotations with known canonical cell type markers (**Fig 2E**), as well as differentially expressed genes between the obtained clusters. Final annotations included 10 different cell types: hematopoietic stem cells (HSCs), megakaryocyte/erythroid (MK_Es), neutrophils (Neutr), B cell progenitors (B_prog), erythroid progenitors (EPs), monocyte progenitors (MPs), megakaryocyte progenitor (MKs), lymphoid-primed multipotent progenitors (LMPPs), basophil, eosinophil and mast cell progenitors (BEMs), and erythroid-megakaryocyte progenitors (EMPs) (**Fig 2F**).

Cell type proportions showed phenotypic heterogeneity between samples, conditions, and datasets overall (**Fig 2F**). Samples coming from the TBD dataset (both TBD patients and healthy donors) seemed to have a higher proportion of HSCs, LMPPs, BEMs and B_prog, while the FA dataset (both FA patients and healthy donors) had more EPs and EMPs. Within the TBD dataset, TBD patients had less LMPPs and HSCs, except for patient TBD_1 which had the highest proportion of HSCs among all samples, and a moderate increase in MK_E compared to healthy controls. On the other hand, between the FA dataset, FA patients had higher proportions of MPs, and less MKs, MK_Es and LMPPs compared to healthy donors. Finally, neutrophiles where a highly variable group among all samples (**Figure 2G**).

### Transcriptional contingency between FA and TBD shows upregulation of inflammatory and cell proliferation pathways

To explore phenotypically relevant commonalities and differences across transcriptional programs in both syndromes, we first calculated differentially expressed genes among healthy, TBD and FA datasets (**Figure 3A**); we then compared the healthy gene expression profile against the TBD & FA (TBD+FA) combined data to highlight transcriptional contingency (**Fig 3B**). Genes sharing downregulation in both syndromes include chromatin remodelers *H1FX* and *H3F3B*, and a group of long non-coding RNAs (*RP11-271C24.2, RP11-1348G14.B, RP11-452H21.4* and *RP11-17M16.2*) with a still poorly known function, and with a bigger reduction in expression in TBD cells (**Fig 3C**).

**Figure 3.**
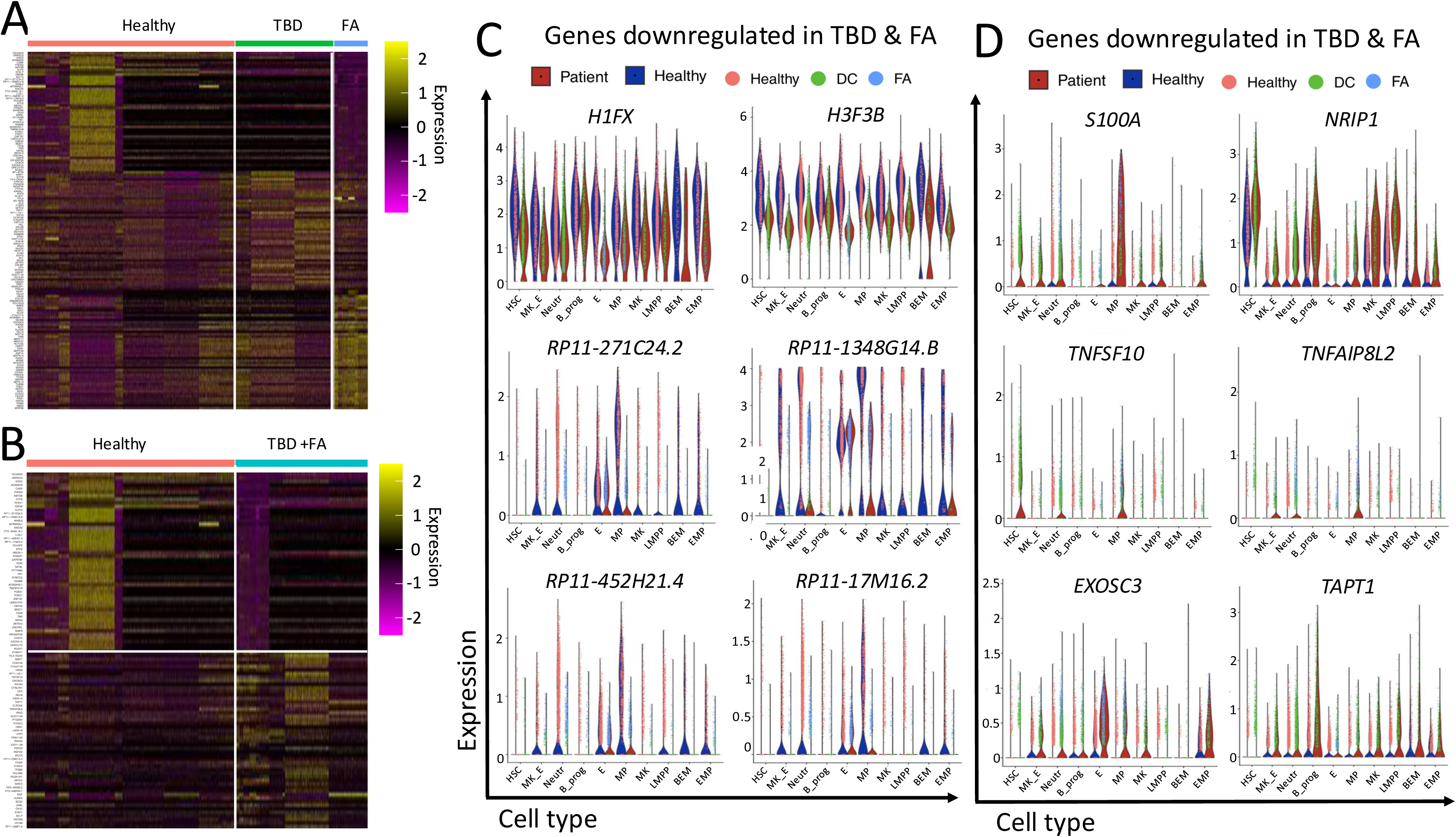
Transcriptional contingency in the HSPCs from FA and DC. **(A)** Heatmap showing the top 100 differentially expressed genes among HSPCs from healthy donors, patients with DC and patients with FA. In this analysis HSPCs from patients with DC and FA were compared as separate groups to obtain differences among each other and with respect to healthy samples. **(B)** Heatmap showing the top 100 differentially expressed genes among HSPCs from healthy donors and patients with DC and patients with FA. In this analysis HSPCs from patients with DC and FA were merged as a single group to obtain transcriptional contingency between DC and FA with respect to healthy samples. **(C)** Violin plots showing genes downregulated in the merged HSPCs from patients with DC and FA, indicating common downregulation. **(D)** Violin plots showing genes upregulated in the merged HSPCs from patients with DC and FA, indicating common upregulation.

On the other hand, upregulated genes in both syndromes included inflammation- related genes like *TNFSF10*, *TNFAIP8L2* and *S100A11*, and proliferation-related genes like *NRIP1*. *TNFSF10* and *TNFAIP8L2* were particularly upregulated in neutrophils, HSCs and MPs (**Figure 3D**). Meanwhile, *S100A11* had a major overall increase in MPs and a preferential increase in EMPs, MKs, MK_Es, LMPPs and HSCs in TBD cells (**Figure 3D**). Finally, other genes with non-related functions were upregulated, such as *NRIP1*, primarily in TBD cells, in B_prog, neutrophiles, MKs, LMPPs and HSCs; *EXOSC3* in Es and EMPs, and *TAPT1* in all cell types but HSCs (**Fig 3D**).

### Inference of differential communication pathways in the HSPC compartment from patients with IBMFS

By pinpointing the transcriptional profiles of the sequenced cells, novel methods for scRNAseq analysis can infer cell-cell communication patterns (31), this is particularly useful in highly heterogeneous samples, such as BM. We took advantage of these strategies and combined the TBD and FA datasets for inferring mechanisms shared by both diseases. This cell communication analysis showed that myeloid progenitors (MP) in the HSPCs from patients with TBD and FA have the highest number of differential interactions (**Figure 4A*, left***), and with a high strength, except for the pairs MP-BEM and MP-LMPP (**Figure 4A*, right***). This suggests that MPs operate important cell-cell communication over other cell types, except for BEMs and LMPPs, in the compartment of HSPCs from patients with IBMFS. Also, LMPP showed decreased values in the differential number and strength of interactions, showing low communication of this cell type with the rest (**Figure 4A and 4B**).

**Figure 4.**
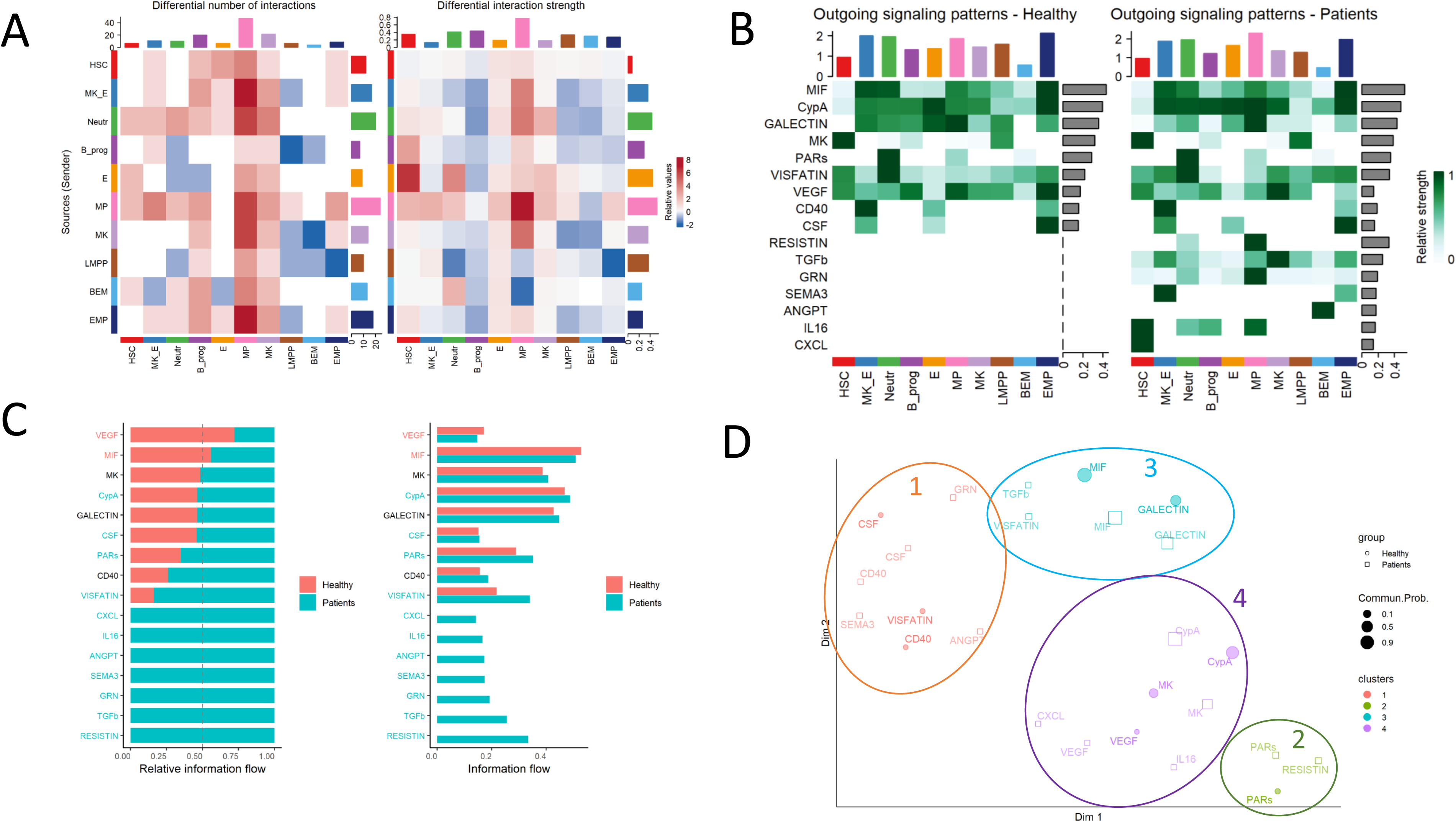
Cell communication in the early HSPCs from IBMFS in comparison to healthy HSPCs. **(A)** Heat map showing cell to cell communication inference. Notably MPs are most connected cells, whereas LMPPs and BEMs show less communications. **(B)** Heat map showing a more complex communication crosstalk in HSPCs from patients. Inflammatory cues are observed in samples from patients. **(C)** Bar plot showing the relative contribution of every pathway to intercellular communication in the HSPCs from DC and FA, in comparison to healthy HSPCs. **(D)** Functional clustering of pathways subtypes enriched in the HSPCs from DC and FA patients with respect to healthy HSPCs.

In addition to inferring communication among cell types, the pathways mediating such communication can be inferred. This analysis shows that, in comparison to the healthy HSPCs, a more complex pattern of communication occurs in the HSPCs from patients with TBD and FA. The cells from IBMFs patients show increased strength in pathways that are seemingly not activated in the healthy HSPCs, including: RESISTIN, TGFβ, GRN, SEMA3, ANGPT, IL16 and CXCL pathways (**Figure 4B**), thus suggesting an increased complexity of the communication network in the BM of these syndromes. Enrichment of pathways being exclusively activated in the HSPCs from TBD and FA cells was further confirmed through information flow analysis, in this analysis, RESISTIN, TGFβ, GRN, SEMA3, ANGPT, IL16 and CXCL pathways were found to be enriched in the cells from patients with IBMFS, whereas the VEGF and MIF pathways is preeminent in the HSPCs from healthy individuals (**Figure 4C**).

Finally, the above-mentioned pathways can be clustered into 4 groups according to their activity type. Cluster 1 is dominated by cell proliferation pathways, cluster 2 by pathway related to inflammation, cluster 3 by immune inhibition pathways and cluster 4 by angiogenesis and migration pathways (**Figure 4D**).

## DISCUSSION

TBDs and FA are the most common and best known IBMFS. These IBMFS share an extremely high risk to develop AA, MDS, AML, and solid tumors, including squamous cell carcinomas (33,34). These striking similarities rise the question whether these syndromes are, at the molecular level, more similar than commonly assumed, and whether the cellular responses to their underlying genetic deficiencies are actually shared.

Some groups have jointly analyzed the BM plasma from multiple IBMFS, including DC and FA, finding increased levels of pro-inflammatory cytokines across the examined IBMFS, and pointing towards shared inflammatory cues in the BM of these patients (37). More recently, the progress in scRNAseq technologies has allowed the obtention of fine-grained transcriptional profiles of HSPCs from patients with IBMFS (35,36,38), allowing the exploration of their transcriptional endeavors, and motivating the search for commonalities and divergences among HSPCs from patients with IBMFS. The shared genomic instability between DC and FA, situates them as a natural starting point for cross-comparisons among IBMFS in the search for their commonalities and divergencies.

As the available scRNAseq datasets from TBD and FA come from different sources (35,36), the first step towards obtaining joint biological conclusions was their integration. However, despite the availability of multiple computational methods, such an integration was not a trivial task. First, it was important to determine which reduced dimensions (PCA) were to be used for integration. For this, a standard analysis (including the determination of the number of variable features to use, normalization, and dimensionality reduction) must have been performed on each of the integrated individual datasets. In this work, we tested methods that are implemented in the Seurat package: FastMNN, scVI, RPCA and Harmony. With the results of each approach, we obtained a *de novo* set of integrated reduced dimensions, which once again had to undergo a standard analysis to determine the best number of variable features and number of dimensions to use for downstream analysis.

To identify the ideal approach, we used the Seurat’s native function MixingMetric that measures the proportion of samples in each cluster/subpopulation. We obtained higher scores with FastMNN and RPCA. To discriminate the approach that better allowed to identify the subpopulations, we visualized the UMAPs created with these two last integration methods. FastMNN better differentiated the clusters of cells, so we decided to use the results yielded by this approach for downstream analysis. In brief, datasets integration is a process that, despite conveniently having multiple computational methods, is very iterative and not straightforward.

After efficient data integration, when we compared the joint gene expression profile of TBD and FA HSPCs, against healthy HSPCs, we observed an aligned downregulation and upregulation of multiple genes. Downregulated genes include the histone genes *H1FX* and *H3FB*. Histones play a critical role in DNA packaging and regulation of gene expression, and their proper function is essential for maintaining genomic integrity. *H1FX* is well-known to be involved in chromatin compaction (21), therefore its reduction, along with *H3FB,* could impair chromatin structure and contribute to the genomic instability seen in these IBMFS, further exacerbating the DNA repair deficiencies characteristic of these syndromes.

Another interesting finding is the concerted downregulation in TBD and FA of a group of poorly characterized long non-coding RNAs (lnc-RNAs), *RP11-271C24.2*, *RP11-1348G14.B*, *RP11-452H21.4* and *RP11-17M16.2*. Lnc-RNAs are emerging as regulators of gene expression, chromatin remodeling, and stem cell function maintenance (22,23). It is therefore tempting to hypothesize that dysregulation of these lnc-RNAs might associate with the disruption of proper HSPCs differentiation and contribute to the impaired hematopoiesis and BM exhaustion observed in FA and TBD.

Concerted upregulation of genes was also observed between TBD and FA, this mainly included inflammation and cell proliferation related genes. *TNFSF10,* for example, is a member of the TNFalpha family of ligands, that has been shown to induce apoptosis in damaged or pre-cancerous cells by engaging their cell death receptors, a mechanism that could be upregulated in IBMFS as a response to chronic DNA damage, and that might thus contribute to HSPCs attrition (24,25). *S100A11,* on the other hand, has been implicated in promoting pro-inflammatory cytokine release, in conditions of sterile inflammation, a condition in which no pathogens are involved (26). This gene expression profile points towards a state of chronic inflammation in TBDs and FA, a well-documented factor that can drive BMF by inducing premature HSPCs exhaustion and that alters the BM niche (27).

Nonetheless, compensatory mechanisms against the growth-suppressive inflammation seem also be taking place, this includes increased expression of proliferative pathways mediated by *NRIP1* (*28*). Interestingly, *NRIP1* has not been previously explored in the IBMFS, and finding its shared upregulation between TBDs and FA makes it an attractive subject of study. Exacerbated proliferative stimuli, are known to promote HSPCs depletion, as most HSPCs enter the cell cycle and terminally differentiate, thus reducing the pool of available long-term HSPCs

The potential role of other upregulated genes in the biology of the IBMFS is less clear, these include *EXOSC3* and *TAPT1*. *EXOSC3* is a core component of the RNA exosome complex, crucial for RNA degradation and processing. Defects in RNA processing could lead to an accumulation of aberrant transcripts, contributing to cellular dysfunction (29). *TAPT1*, in turn, is involved in skeletal development, and has been linked to ciliogenesis and cell signaling pathways. While its role in BM biology is unclear, altered signaling or ciliary dysfunction could impact stem cell niche interactions or intracellular signaling pathways crucial for hematopoiesis (30). Further study of these genes in the context of FA and TBD is warranted to elucidate their potential contributions to BMF.

Finally, inference of cell-cell communication cues further suggests that strong inflammatory signals mediate the communication among HSPCs in patients with IBMFS. The relevance of the TGFβ pathway and IL-16, two well-known growth suppressive pathways, as mediators of cell-cell communication was confirmed in these cells and seems to be a common theme between FA and TBDs. Interestingly, the cells with the highest outgoing signaling patterns, and towards other cell types, are myeloid progenitors, whether this increased communication is related to the myeloid-biased differentiation of HSPCs that is commonly observed in patients with IBMFS or related to their increased risk to develop myeloid malignancy is a matter that deserves further elucidation.

## CONCLUSIONS

In this work we have successfully integrated publicly available scRNAseq datasets of HSPCs from patients with TBDs and FA, the most common IBMFS. Analysis of these integrated HSPCs shows commonalities and differences between these syndromes, including concerted downregulation and upregulation of subsets of genes. An important commonality includes a strong inflammatory signal seemingly mediated by the TGFβ pathway and by members of the TNFα family of ligands.

## ACKNOWLEDGENTS

This research was funded by the “Programa de Apoyo a Proyectos de Investigación e Innovación tecnológica (PAPIIT) through the project IA205022” and the “Consejo Nacional de Humanidades, Ciencias y Tecnologías (CONAHCyT) through the project 319344”. PS receives the CONAHCyT scholarship CVU 2007450. PLA receives the CONAHCyT scholarship CVU 941552.

## Data Availability Statement

The integrated and harmonized datasets are preliminarily available at: https://drive.google.com/file/d/1zKT0FdWb02-chdt9iHbyONABHzcic0-5/view?usp=drive_link

